# Benchmarking joint multi-omics dimensionality reduction approaches for cancer study

**DOI:** 10.1101/2020.01.14.905760

**Authors:** Laura Cantini, Pooya Zakeri, Celine Hernandez, Aurelien Naldi, Denis Thieffry, Elisabeth Remy, Anaïs Baudot

**Author notes:** Centre for Brain and Disease Research, Flanders Institute for Biotechnology (VIB), Leuven, Belgium and Department of Neurosciences and Leuven Brain Institute, KU Leuven, Leuven, Belgium. Institute for Integrative Biology of the Cell (I2BC), CEA, CNRS, Univ. Paris-Sud, Université Paris-Saclay, 91198, Gif-sur-Yvette cedex, France.

## Abstract

High-dimensional multi-omics data are now standard in biology. They can greatly enhance our understanding of biological systems when effectively integrated. To achieve this multi-omics data integration, Joint Dimensionality Reduction (jDR) methods are among the most efficient approaches. However, several jDR methods are available, urging the need for a comprehensive benchmark with practical guidelines.

We performed a systematic evaluation of nine representative jDR methods using three complementary benchmarks. First, we evaluated their performances in retrieving ground-truth sample clustering from simulated multi-omics datasets. Second, we used TCGA cancer data to assess their strengths in predicting survival, clinical annotations and known pathways/biological processes. Finally, we assessed their classification of multi-omics single-cell data.

From these in-depth comparisons, we observed that intNMF performs best in clustering, while MCIA offers a consistent and effective behavior across many contexts. The full code of this benchmark is implemented in a Jupyter notebook - multi-omics mix (momix) - to foster reproducibility, and support data producers, users and future developers.

## Background

Due to the advent of high-throughput technologies, high-dimensional “omics” data are produced at an increasing pace. In cancer biology, in particular, national and international consortia, such as The Cancer Genome Atlas (TCGA), have profiled thousands of tumor samples for multiple molecular assays, including mRNA, microRNAs, DNA methylation and proteomics ^1^. Moreover, multi-omics approaches are currently being transposed at single-cell level, which further stresses the need for methods and tools enabling the joint analysis of such large and diverse datasets ^2^.

While multi-omics data are becoming more accessible, and studies combining different omics more frequent, the genuine joint analysis of multi-omics data remain very rare. Achieving proper multi-omics integration is crucial to bridge the gap between the vast amount of available omics and our current understanding of biology. By integrating multiple sources of omics data, we can reduce the effect of experimental and biological noise. In addition, different omics technologies are expected to capture different aspects of cellular functioning. Indeed, the different omics are complementary, each omics containing information that is not present in others, and multi-omics integration is thereby expected to provide a more comprehensive overview of the biological system. In cancer research, omics have been profiled at different molecular layers, such as genomics, transcriptome, epigenome, and proteome. Integrating these large-scale and heterogeneous sources of data allows researchers to address crucial objectives, including (i) classifying cancer samples into subtypes, (ii) predicting the survival and therapeutic outcome of these subtypes, and (iii) understanding the underlying molecular mechanisms that span through different molecular layers ^3^.

Designing theoretical and computational approaches for the joint analysis (aka intermediate integration) of multi-omics datasets is currently one of the most relevant and challenging questions in computational biology ^3,4^. Indeed, the different types of omics have a large number of heterogeneous biological variables and a relatively low number of biological samples, thereby opening to all the challenges typical of “Big Data”. In addition, each omics has its own technological limits, noise, and range of variability. All these elements can mask the underlying biological signals. Multi-omics integrative approaches should be able to capture not only signals shared by all omics data but also those emerging from the complementarity of the various omics data.

The joint analysis of multiple omics can be performed with various integrative approaches, classified in broad categories ^4,5^. Bayesian methods, such as Bayesian Consensus Clustering (BCC) ^6^, build a statistical model by making assumptions on data distribution and dependencies. Network-based methods, such as Similarity Network Fusion (SNF) ^7^, infer relations between samples or features in each omics layer, and further combine the resulting networks. Dimensionality Reduction (DR) approaches decompose the omics into a shared low-dimensional latent space ^8,9^. Four recent reviews tested and discussed some of these existing methods from the clustering performance perspective ^10–13^. Pierre-Jean et al. ^12^, Rappoport et al.^10^ and Tini et al.^13^ selected one method from each of the aforementioned three categories, while Chauvel et al.^11^ focused on Bayesian and DR approaches.

From these initial reviews, DR approaches emerged as particularly well-performing. They are well-adapted to solve high-dimensional mathematical problems. Furthermore, the richness of the information contained in their output enhances their relevance for multi-omics integration. Indeed, DR methods enable the classification of samples (clustering/subtyping), the clinical characterization of the identified clusters/subtypes and a variety of other downstream analyses, including the analysis of cellular processes and/or pathways (Figure 1). Thus, DR simultaneously provides information on all the key objectives mentioned above, namely the classification of samples into subtypes, their association with outcome/survival as well as the reconstruction of their underlying molecular mechanisms. As a consequence, the design of DR approaches for the joint analysis and integration of multiple omics (jDR) is currently a highly active area of research ^5,8,10,11,14^.

**Figure 1.**
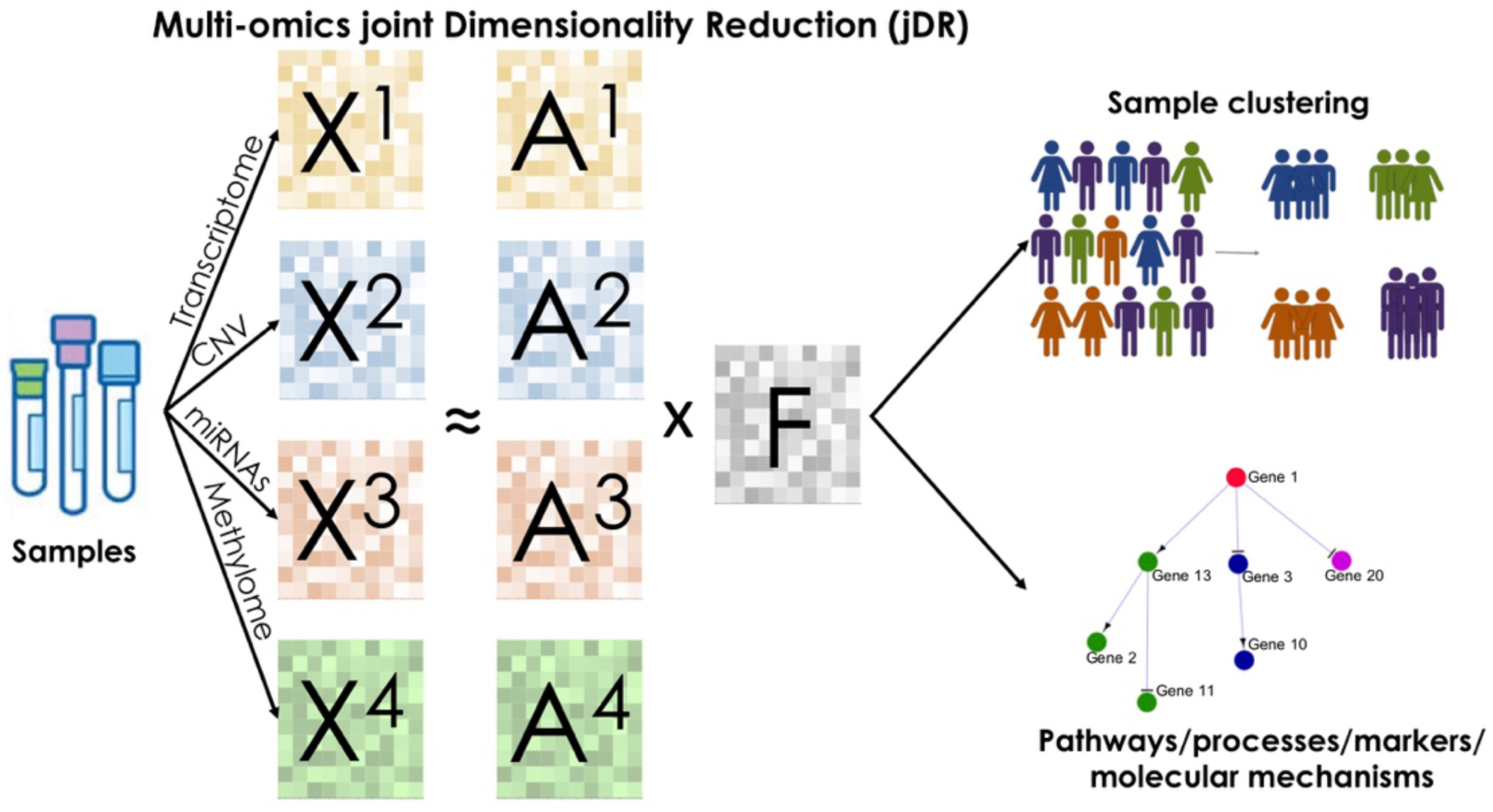
Joint Dimensionality Reduction methods overview. Multi-omics are profiled from the same set of samples. Each omics corresponds to a different matrix *X*^*i*^. jDR methods factorize the matrices X^i^ into the product of a factor matrix *F* and weight matrices *A*^*i*^. These matrices can then be used to cluster samples and identify molecular processes.

Here, we report an in-depth comparison of representative state-of-the-art multi-omics joint Dimension Reduction (jDR) approaches, in the context of cancer data analysis. We extensively benchmarked nine approaches, spanning the main mathematical formulations of multi-omics jDR, in three different contexts. First, we simulated multi-omics datasets and evaluated the performance of the nine jDR approaches in retrieving ground-truth sample clustering. Second, we used TCGA multi-omics cancer data to assess the strengths of jDR methods in predicting survival, clinical annotations, and known pathways/biological processes. Finally, we evaluated the performance of the methods in classifying multi-omics single-cell data from cancer cell lines.

All these analyses allow formulating recommendations and guidelines for users, as well as indications for methodological improvements for developers. We also provide the Jupyter notebook multi-omics mix (momix) and its associated Conda environment containing all the required libraries installed (https://github.com/ComputationalSystemsBiology/momix-notebook). Overall, momix can be used to reproduce the benchmark, but also to test jDR algorithms on other datasets, and to evaluate novel jDR methods and compare them to reference ones.

## Results

### 1. Joint Dimensionality Reduction approaches and principles

The goal of joint Dimensionality Reduction (jDR) approaches is to reduce high-dimensional omics data into a lower dimensional space. We consider *P* omics matrices *X*^*i*^, *i* = 1, …, *P* of dimension *n* × *m*, with *n* features (e.g. genes, proteins) and *m* samples. A jDR jointly decomposes the P omics matrices into the product of a *k* × *m factor matrix (F*)and *n* × *k* omics-specific *weight/projection matrices (A*^*i*^) (Figure 1). Here and in the following, we will denote as *factors* the columns of the factor matrix and as *metagenes* the rows of the weight/projection matrix corresponding to transcriptomic data (Methods). A description of the mathematical formulations of the nine jDR approaches is provided in the Method section.

A wide variety of methods exist to perform jDR (Supplementary Table 1). These methods are based on different underlying mathematical formulations, including Principal Components Analysis, Factor analysis, co-inertia analysis, Gaussian latent model, matrix tri-factorization, Non-negative Matrix Factorization, CCA or tensor representations (Supplementary Table 1). We selected nine jDR approaches representative of each of these main mathematical formulations (Table 1), focusing on methods able to combine more than two omics, implemented in R or Python, and with software readily available and documented. These jDR approaches are iCluster^15^, Integrative NMF (intNMF)^16^, Joint and Individual Variation Explained (JIVE)^17^, Multiple co-inertia analysis (MCIA)^18^, Multi-Omics Factor Analysis (MOFA)^19^, Multi-Study Factor Analysis (MSFA)^20^, Regularized Generalized Canonical Correlation Analysis (RGCCA)^21^, matrix-tri-factorization (scikit-fusion)^22^ and tensorial Independent Component Analysis (tICA)^23^.

**Table 1.**
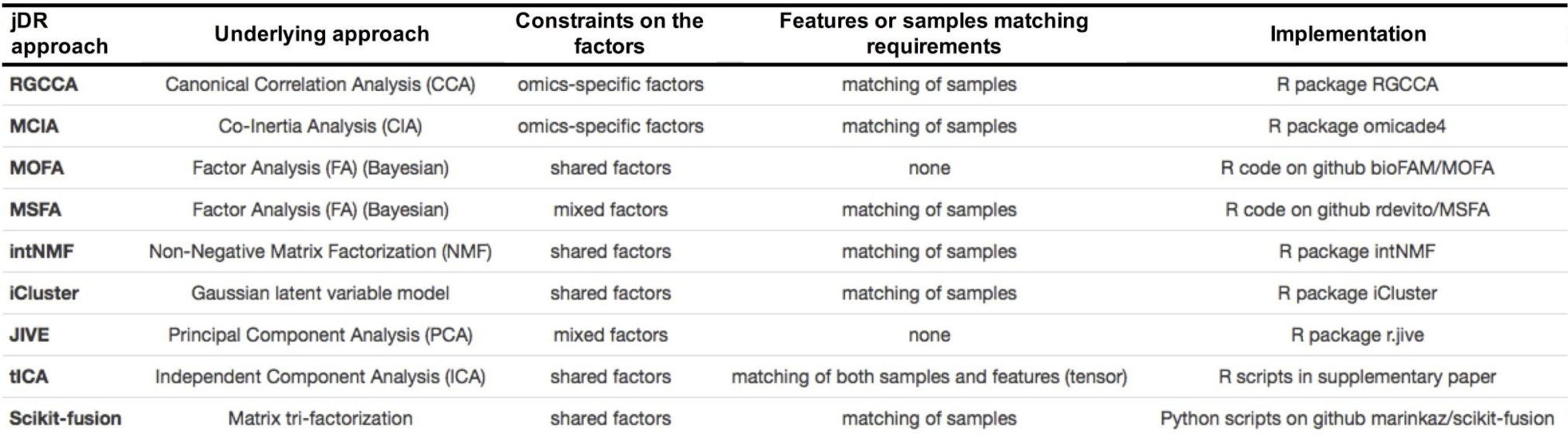
Selected nine joint Dimensionality Reduction approaches benchmarked in this study.

Some of the selected nine jDR approaches are extensions of DR methods initially designed for single omics datasets: intNMF is an extension of non-Negative Matrix Factorization (NMF); tICA is an extension of Independent Component Analysis (ICA); MCIA and JIVE are different extensions of Principal Component Analysis (PCA); and MOFA, MSFA, and iCluster are extensions of Factor Analysis. As a consequence, the different jDR algorithms make different assumptions on the distribution of the factors (Methods). The different jDR approaches also make different assumptions on the across-omics constraints on the factors. Some algorithms, such as MOFA, consider the factors to be *shared* across all omics datasets. In contrast, the factors of RGCCA and MCIA are different for each omics layer, i.e., they are *omics-specific* factors. These omics-specific approaches still maximize some measures of interrelation between the omics-specific factors such as their correlation (RGCCA), or their co-inertia (MCIA). Finally, JIVE and MSFA consider *mixed* factors, decomposing the omics data as the sum of two factorizations, one containing a unique factor matrix common to all omics, and the second having omics-specific factor matrices. These last two categories of methods, omics-specific and mixed thus address also the complementarity of the multi-omics.

The majority of the jDR approaches can manage different features (e.g., genes, miRNAs, CpGs…), but require a match between the samples of the different omics datasets (columns of the *X*^*i*^matrices, see Table 1). Some algorithms, such as MOFA, scikit-fusion and JIVE, can also cope with omics matrices having not all samples in common. This is particularly suitable for multi-omics integration given that missing samples are frequent in data collections, such as in TCGA. For the sake of comparison, we applied here all methods considering only the samples profiled for all omics. Tensorial approaches, represented by tICA, require by definition that all matrices*X*^*i*^ have exactly the same samples and features. Nonetheless, the features of multi-omics data are frequently different (e.g. genes, miRNAs). A possible strategy to have the same features for all omics would be to convert all the features to the same level, e.g. gene symbols. This is sometimes unfeasible: miRNAs cannot be converted to gene symbols, for instance. We applied here another strategy, where we considered for each omics the matrix of correlation-of-correlation between samples (Methods). Both strategies imply a loss of information, which can affect the results of the omics integration.

Noteworthy, intNMF and iCluster produce, in addition to the factors, a clustering of the samples. Scikit-fusion can combine omics data with additional annotation (i.e., side information, such as pathway or process annotations). However, for the sake of comparisons with other algorithms, scikit-fusion is applied here without side-information.

### 2) Benchmarking joint Dimensionality Reduction approaches on simulated omics datasets

We first evaluated the jDR approaches on artificial multi-omics datasets. We simulated these datasets using the *InterSIM* CRAN package ^24^. This package generates three artificial omics datasets with imposed reference clustering by manipulating TCGA ovarian cancer data. Thereby, it avoids making assumptions on the distribution of the simulated data. We simulated multi-omics data with five, ten, and fifteen clusters. In addition, each set of clusters is simulated in two versions, either with all clusters of the same size, or with clusters of variable random sizes (Methods).

We applied the nine jDR methods, requiring the decomposition of multi-omics data into five, ten, and fifteen factors, depending on the simulated datasets. The performances of the nine jDR approaches are then compared based on their clustering of samples. As mentioned before, intNMF and iCluster are intrinsically designed for sample clustering, while the remaining seven algorithms detect factors without providing a direct clustering. Accordingly, we applied directly intNMF and icluster. For the seven other algorithms, we obtained the clustering of the samples by applying consensus clustering to the factor matrix (Methods) ^25^.

The agreement between the clustering obtained by the various jDR algorithms and the ground-truth clustering is measured with the Jaccard Index (JI) (Methods). First, we observed that all methods perform reasonably well in the different simulated scenarios (JIs >= 0.6, Figure 2). The two algorithms intrinsically designed for clustering, namely intNMF and iCluster, display the best performances. In particular, intNMF retrieves perfectly the ground-truth clusters (JI ∼ 1). iCluster presents some variability for five and ten clusters, independently of the size distribution of the clusters. Regarding the remaining seven jDR approaches, MCIA, MOFA, and RGCCA are overall the best-performing methods. These methods are indeed among the top-three best algorithms in 6/6, 6/6, and 5/6 simulated scenarios, respectively. tICA and scikit-fusion are the less effective methods in this benchmark. tICA structures the multi-omics data into a tensor. As described previously, to obtain these tensors, we transformed the omics data into correlation-of-correlation matrices, which might induce a loss of information. scikit-fusion is designed to work with side information, which are used to build a relation network connecting the various entities (e.g. samples, genes, proteins). However, for the sake of comparison with the other jDR methods, side-information was not considered, and this could have affected the results of the algorithm.

**Figure 2.**
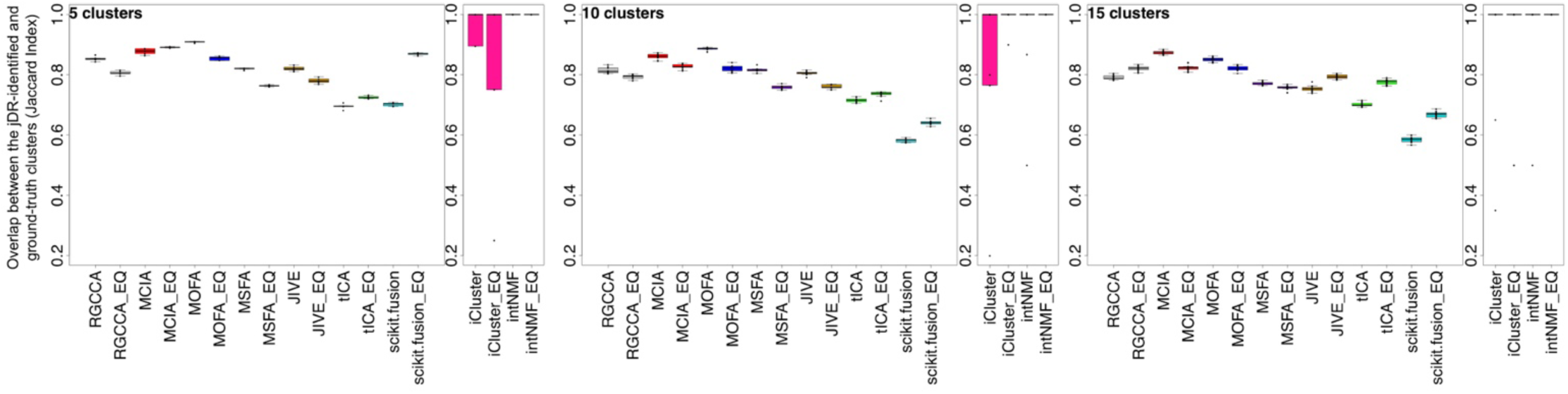
jDR clustering of simulated multi-omics datasets. Boxplots of the Jaccard Index computed between the clusters identified by the different jDR methods and the ground-truth clusters imposed on the simulated data (for 5, 10 and 15 imposed clusters). For each method (e.g. RGCCA), performances on heterogeneous and equally-sized clusters are reported (denoted as RGCCA and RGCCA_EQ, respectively).

### 3) Benchmarking joint Dimensionality Reduction approaches on cancer datasets

In the second step, we downloaded TCGA multi-omics data for ten different cancer types ^10^ (https://portal.gdc.cancer.gov/). These data are composed of three omics layers: gene expression, DNA methylation, and miRNA expression. The number of samples ranges from 170 for Acute Myeloid Leukemia (AML) to 621 for Breast cancer. We applied the nine jDR approaches to each of these cancer multi-omics datasets, jointly decomposing them in ten factors, as in the work of Bismeijer and colleagues ^26^. Most cancer subtyping approaches indeed revealed ten or fewer clusters of samples (i.e., subtypes). The Factor Analysis approach MSFA did not converge to any solution and was thereby not further considered. Importantly, we do not have ground-truth for cancer subtyping (i.e. clustering of cancer samples). We hence compared the performances of the remaining eight jDR approaches regarding their ability to identify factors predictive of survival, as well as factors associated with clinical annotations. We also evaluated the weight matrices resulting from the jDR methods, by assessing their enrichment in known biological pathways and processes.

To test the association of the jDR factors with survival, we used the Cox proportional-hazards regression model. We observed first that the number of factors associated with survival depends more on the cancer types than on the jDR algorithm (Figure 3 and Supplementary Figure 1). Indeed, for three cancer types (Colon, Lung, and Ovarian), none of the jDR methods was able to identify survival-associated factors. This result is in agreement with previous observations testing the association of multi-omics clusters with survival on the same TCGA data with the log-rank test ^10^. In four other cancer types (Sarcoma, Liver, Kidney, and Breast), all jDR algorithms identified one or two survival-associated factors. Finally, in Melanoma, GBM, and AML, the majority of the jDR methods identified three or four survival-associated factors. In general, MCIA, RGCCA, and JIVE achieved the best performances, finding factors significantly associated with survival in seven out of ten cancer types. These approaches also offered the most significant p-values in the higher number of cancer types. JIVE is the best approach in three cancer types: Liver cancer (p-value ∼ 10^−4^), AML (10^−3^) and in GBM (10^−3^); RGCCA is the best in Melanoma (10^−8^) and Breast cancer (10^−3^); and MCIA is the best in Kidney cancer (10^−4^) and Sarcoma (10^−5^). Furthermore, RGCCA, MCIA, and JIVE showed the most promising results for the cancer types having overall less survival-associated factors (Sarcoma, Liver, Kidney, and Breast, Figure 3 and Supplementary Figure 1).

**Figure 3.**
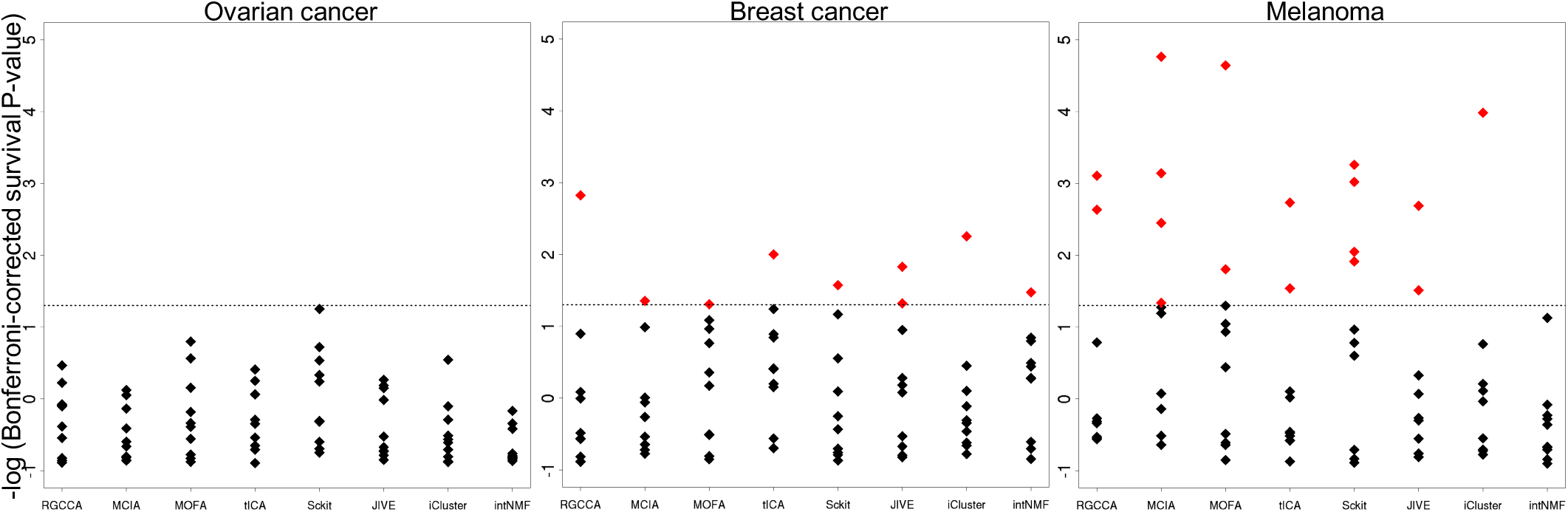
Identification of factors predictive of survival in Ovarian, Breast and Melanoma cancer samples by the jDR methods. For each method the Bonferroni-corrected p-values associating each of the 10 factors to survival (Cox regression-based survival analysis) are reported. The dot lines correspond to a corrected p-value threshold of 0.05. The results corresponding to the other seven cancer types are presented in Supplementary Figure 1.

Afterward, we assessed the association of the jDR factors with clinical annotations. We selected four clinical annotations: “age of patients,” “days to new tumor,” “gender”, and “neo-adjuvant therapy somministration” (Methods). To test the significance of the associations of the factors identified by the jDR methods with these clinical annotations, we used Kruskal-Wallis tests for multi-class annotations (“age of patients” and “days to new tumor”), and Wilcoxon rank-sum for binary annotations (“gender” and “neo-adjuvant therapy somministration”). In addition, we intended to evaluate the methods not only by their capacity to associate factors with clinical annotations, but also by their ability to achieve these associations with a one-to-one mapping between a factor and a clinical annotation, i.e. their selectivity (Figure 4). Indeed, a jDR method detecting one factor associated with multiple clinical annotations cannot distinguish the annotations from each other. To the contrary, a jDR method detecting multiple factors associated with only one clinical annotation does not maximally explore the spectrum of all possible annotations. We defined a selectivity score having a maximum value of 1 when each factor is associated with one and only one clinical annotation, and a minimum of 0 when all factors are associated with all clinical annotations (Methods). The average selectivity value of all methods across all cancer types is 0.49. The top methods in each cancer type are defined as those having a maximum number of factors associated with clinical annotations, together with a selectivity value above the average. RGCCA, MCIA, and MOFA are overall the best-performing algorithms, since they rank among the top three methods in 6/10, 5/10, and 5/10 cancer types, respectively. In contrast, intNMF, scikit-fusion, and tICA are less effective (among the top three methods in only two out of ten cancer types).

**Figure 4.**
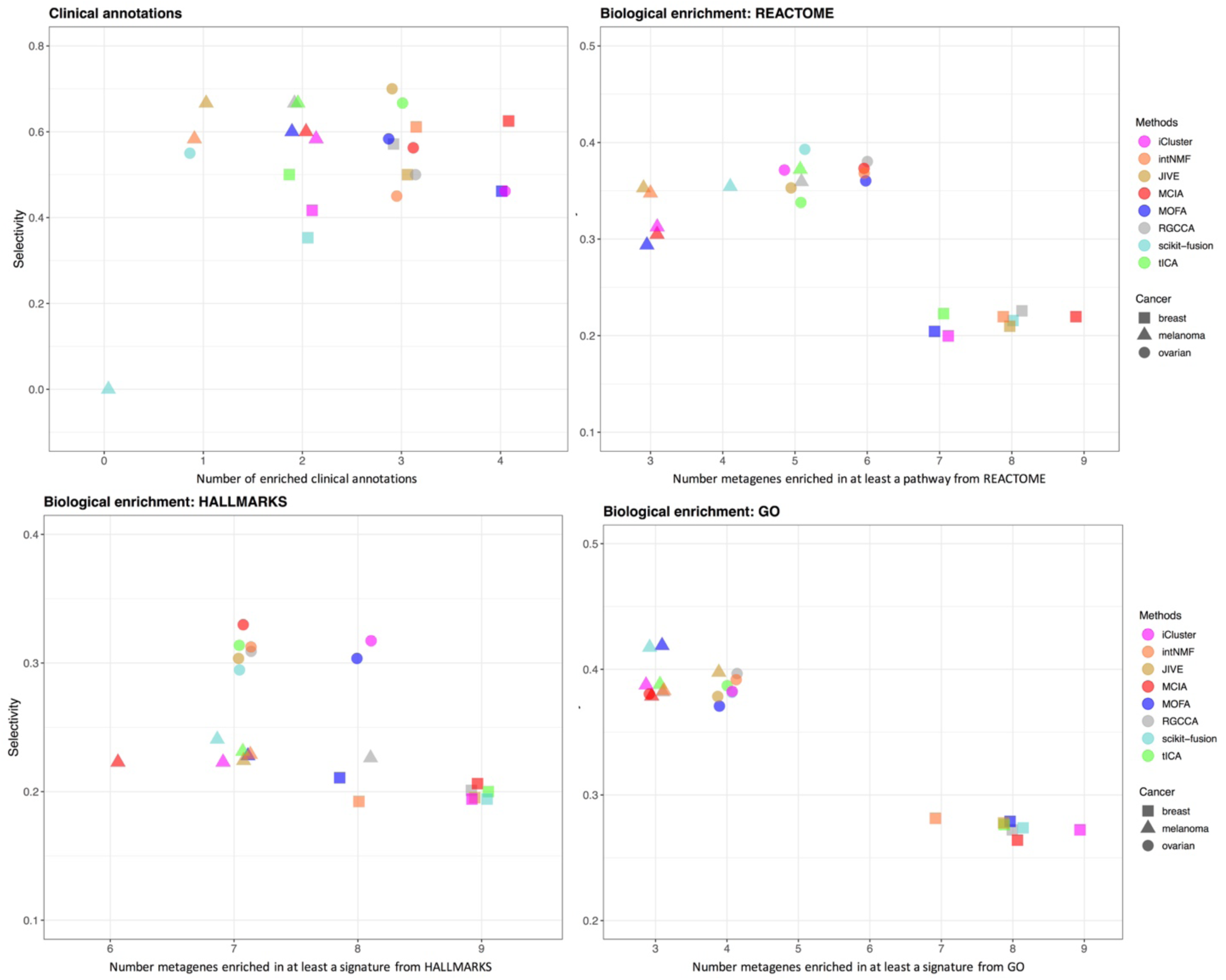
Identification of factors associated with clinical annotations, and metagenes associated with biological annotations in Ovarian, Breast and Melanoma samples, by the jDR methods. For clinical annotations, the plot represents, for each method, the number of clinical annotations enriched in at least one factor together with the selectivity of the associations between the factors and the clinical annotations (Method). For the three annotation sources (MsigDB Hallmarks, REACTOME and Gene Ontology), the number of metagenes identified by the different jDR methods enriched in at least a biological annotation are plotted against the selectivity of the associations between the metagene and the annotation. See Supplementary Figure 2 for the results corresponding to the other seven cancer types.

Finally, we assessed the jDR methods performances in associating factors with biological processes and pathways (Figure 4). To achieve this goal, we need to take into account genes (i.e., weight matrices) and not samples (i.e., factor matrices). We computed the number of metagenes (corresponding to the rows of the transcriptomics weight matrix) enriched in at least one biological annotation from Reactome, Gene Ontology (GO), and cancer Hallmarks annotation databases (Methods). An optimal jDR method should maximize the number of metagenes enriched in at least one biological annotation, while optimizing also the selectivity (defined as above for clinical annotations and in the Methods). The average selectivity of all methods across the ten cancers is 0.3 for Reactome, 0.35 for GO, and 0.26 for cancer Hallmarks. The top methods in each cancer type are defined as those having a maximum number of metagenes associated with biological annotations, together with selectivity values above the average. Scikit-fusion, tICA, and RGCCA are overall the best-performing algorithms for Reactome annotations (ranking among the top three methods in 4/10, 3/10, 3/10 cancer types, respectively). tICA, iCluster and MCIA offered the best performances in cancer Hallmarks annotations (ranked among the top three methods in 4/10, 3/10, 3/10 cancers, respectively) and MCIA, intNMF and iCluster performed the best in GO annotations (ranked among the top three methods in 4/10, 3/10, 3/10 cancers, respectively). Overall, among all jDR methods, tICA and MCIA offer the most promising results for two out of three annotation databases considered in this study and they get the best average performances across the three annotations databases (Table 1).

### 4) Benchmarking joint Dimensionality Reduction approaches on single-cell datasets

Similarly to bulk multi-omics analyses, the joint analysis of single-cell multi-omics is expected to provide tremendous power to untangle the cellular complexity. In addition, jDR approaches could compensate for the strong intrinsic limitations of single-cell multi-omics, such as small number of sequencing reads, systematic noise due to the stochasticity of gene expression at single-cell level, or dropouts. However, the nine jDR algorithms that we are considering (excepted MOFA) have been designed and applied to bulk multi-omics data. It is therefore crucial to evaluate and benchmark the performances of these jDR algorithms for single-cell multi-omics integration.

To test the jDR approaches on single-cell datasets, we fetched scRNA-seq and scATAC-seq, simultaneously measuring gene expression and chromatin accessibility on three cancer cell lines (HTC, Hela and K562) for a total of 206 cells, and reported in the study of Liu and colleagues ^27^. As these cells have been obtained from three different cancer cell lines, we expect that the first two factors of the various jDR approaches would cluster single-cells according to their cancer cell line of origin.

The first two factors of the nine jDR algorithms show overall good performances to cluster cells according to cell lines of origin (Figure 5). To compare quantitatively these clustering results, we measured the C-index with values in the range [0,1], where 0 represents an optimal clustering. According to our results, tICA and MSFA are best-performing jDR methods with a C-index of 0, immediately followed by MCIA and intNMF (C-indices 0.018 and 0.025, respectively), followed by RGCCA, MOFA, and scikit-fusion(C-indices 0.077, 0.12, 0.19, respectively), and finally, JIVE and iCluster (C-indices 0.23 and 0.25, respectively).

**Figure 5.**
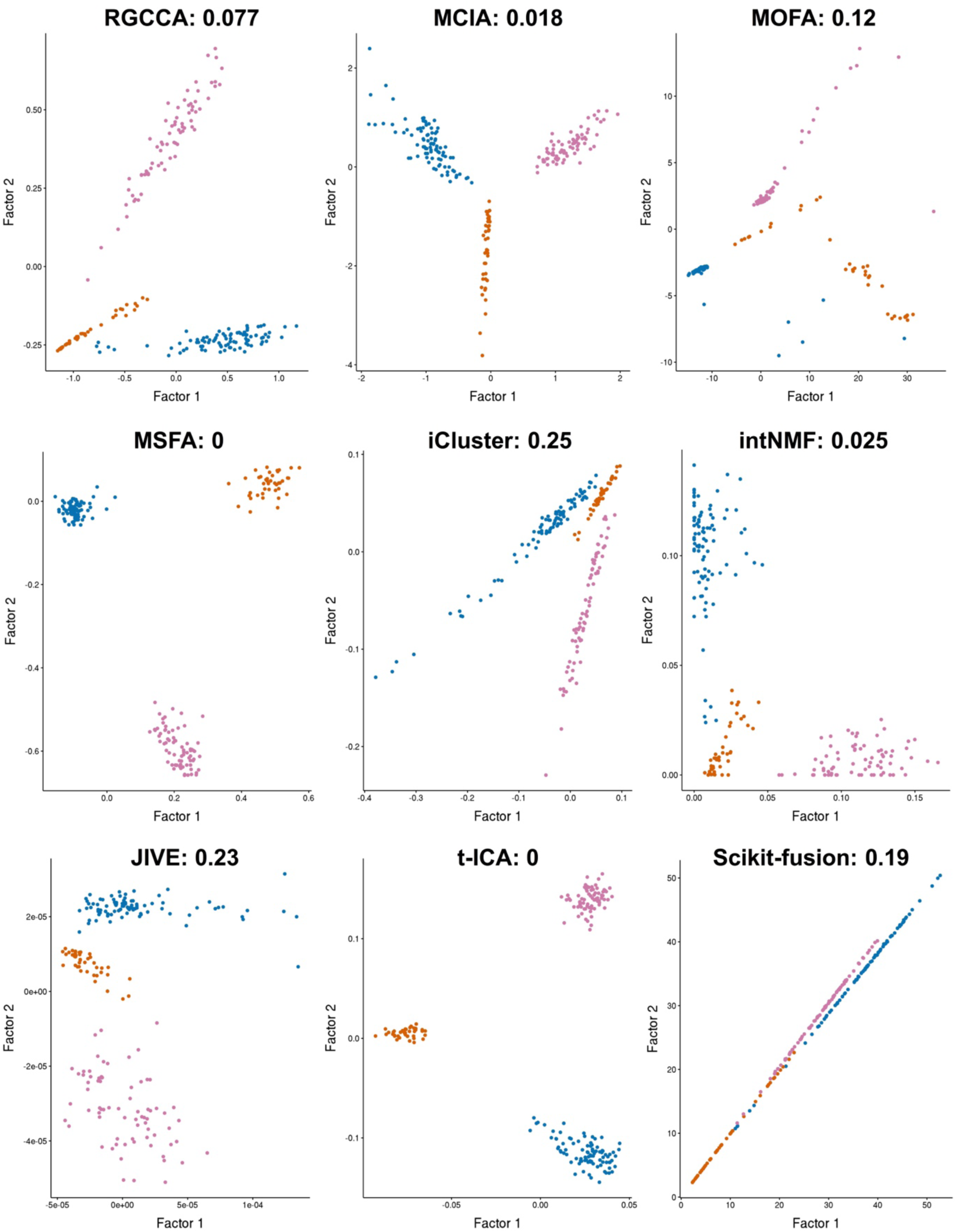
jDR clustering of single-cell multi-omics according to the cancer cell line of origin. Scatterplots of factor 1 and 2 (i.e., the first two columns of the factor matrix) are reported for each jDR method. The colors denote the cancer cell line of origin: pink for K562, orange for Hela and blue for HCT. The C-index (in the range [0-1]) reports the quality of the obtained clusters (0 being the best).

### 5) Multi-omics mix (momix) Jupyter notebook

To foster the reproducibility of all the results and figures presented in this benchmark study, we implemented the corresponding code in a Jupyter notebook available at https://github.com/ComputationalSystemsBiology/momix-notebook together with the associated Conda environment containing all the required libraries installed. Written in R, this notebook is structured in three main parts corresponding to the three test cases here considered (simulated data, bulk TCGA cancer data and single-cell data). Importantly, this notebook can be easily modified to test the nine jDR algorithms on user-provided datasets. The notebook can also be adjusted to benchmark novel jDR algorithms on our three test cases. Full documentation to achieve these goals is included in the notebook.

## Discussion

We benchmarked in-depth nine jDR algorithms, representative of multi-omics integration approaches, in the context of cancer data analysis. In contrast to existing comparisons ^10–13^, our benchmark not only focuses on the evaluation of the clustering outputs, but also evaluates the biological, clinical, and survival annotations of the factors and metagenes. Existing comparisons also mainly use simulated data; we here consider large datasets of bulk cancer multi-omics as well as single-cell data.

When performing clustering on simulated multi-omics datasets, intNMF, intrinsically designed as a clustering algorithm, offered the most promising results. In the same sub-benchmark, MCIA, MOFA, and RGCCA showed the best performance among the set of methods not intrinsically designed for clustering. In the cancer data sub-benchmark, when we evaluated the associations of the factors with survival or clinical annotations, MCIA, JIVE, MOFA, and RGCCA were the most efficient methods. When assessing the associations of the metagenes with biological annotations, MCIA and tICA were the most efficient. Finally, in the last sub-benchmark, when clustering single-cell multi-omics data, MSFA and tICA, as well as MCIA and intNMF, outperformed other approaches.

As mentioned earlier, intNMF, representative of the Non-negative Matrix Factorization (NMF) approaches, performs well for the clustering tasks, i.e., for detecting substantial patterns of variation across the omics datasets. This is observed for both simulated bulk data clustering and single-cell data clustering. Hence, intNMF should be prioritized by researchers focusing on clustering samples. However, intNMF is not effective when assessing the quality of individual factors and metagenes, as observed in the bulk cancer sub-benchmark. Our results rather suggest that researchers interested in exploring factor-level information, such as associations with clinical annotation or survival, should rather consider MCIA, JIVE, MOFA and RGGCA. When focusing on the underlying biology of the metagenes, tICA and MCIA should be prioritized. Indeed, we showed that these approaches are efficient to detect pathways or processes, but they could also be interesting to identify biomarkers or other molecular mechanisms. Finally, our study highlights the potentiality of MCIA. Indeed, MCIA is the method with the most consistent and effective behavior across all the different sub-benchmarks. It can thereby be useful for researchers interested in applying jDR without favouring any particular biological question.

In the future, it would be interesting to extend our benchmark to evaluate the jDR methods also integrating discrete omics data. Indeed our current benchmark focuses on continuous data (e.g. expression, methylation), whereas many -omics and annotations can be formalized as discrete data (e.g. copy number, mutation, drug response). Further extensions of our benchmark could also investigate the impact of different variables on the jDR methods, such as the stability of the methods with respect to variations in the structure of omics data (e.g. imbalance in variability or number of features); or optimal performances according to different combinations of omics data (e.g. are three omics more informative than two?). In addition, to make a fairer comparison, we imposed the same numbers of factors to all of jDR methods, but we could imagine using the optimal number of factors directly computed by each method, as in the work of Tini and colleagues ^13^. Finally, multi-omics data are frequently profiled from different sets of patients/samples, leading to missing data, and further extensions of the benchmark could take this point into account.

Among the methods selected in our benchmark, MOFA is the only approach already tested for the multi-omics integration of single-cell data. But recently, other jDR methods have been published for this purpose: LIGER ^28^, Seurat ^29^ and MOFA+ ^14^, for instance. However, these single-cell jDR approaches cannot be evaluated on our first two sub-benchmarks that focus on bulk data. In our single-cell sub-benchmark, we evaluated the jDR approaches for their clustering capacities. This evaluation should be complemented in the future by benchmarks focusing on single-cell multi-omics integration, and retrieving also pseudo-temporal trajectories, for instance.

From a technical perspective, we observed that the methods that seek for omics-specific factors often led to a better performance than the methods designed for finding shared or mixed factors. We hypothesize that jDRs with omics-specific factors could successfully detect not only biological processes shared across multiple omics but also those processes that are complementary in multiple sources of omics data. In addition, when using algorithms having omics-specific factors, we only evaluated the transcriptome-associated omics-specific factors (Methods). The outputs of these methods often contain relevant information, such as additional omics-specific factors. The use of co-inertia (as implemented in MCIA) further seems more efficient to enforce relationships across omics than the use of correlation (as implemented in RGCCA). Accordingly, we suggest developers to prioritize omics-specific factors for further methodological developments. In addition, there is room for development of approaches managing missing data, as many of the best performing approaches, such as MCIA, can work only on omics profiled from the same samples. This is also true for the consideration of discrete data as among the methods considered here, only MOFA and scikit-fusion have been previously applied to such data. Finally, most of the considered methods detect only linear signals. MOFA, in particular in its most recent version (MOFA+)^14^, is the only algorithm in our benchmark that can also detect slightly nonlinear signal. As a result, future developments should be directed towards methods that can capture the nonlinear signals present in the data. Developers could take advantage of the momix Jupyter notebook using it to compare novel methods with established ones.

## Methods

We consider P omics matrices *X*^*i*^, *i* = 1, . ., *P*, each of dimension (*n* × *m*), where the n lines correspond to the features (e.g. genes, miRNAs, CpGs), and the m columns correspond to the samples. jDR jointly decomposes *X*^*i*^, *i* = 1, . ., *P*, into a factor matrix (of dimension *k* × *m*) and omics-specific weight matrices (of dimension *n* × *k*). We will denote as *factors* the columns of the factor matrix and *metagenes* the rows of the weight matrix associated with transcriptome.

### Presentation of the nine jDR algorithms

We detail here the nine jDR methods benchmarked in momix. We selected default parameters for each approach. Each method can in principle optimize its number of factors to be detected, but for the sake of comparison, we imposed the same number of factors on all approaches. Please note that we followed the mathematical formulations and notations provided in each method publication.

#### 1. Integrative Non-negative Matrix Factorization (intNMF)

intNMF ^16^ is one of the numerous generalizations of NMF to multi-omics data. The method decomposes each omics matrix *X*^*i*^ into a product of non-negative matrices: the factor matrix W, and an omics-specific matrix *H*^*i*^

*X*^*i*^ = *WH*^*i*^, for *i* = 1. …, *P* with *W* and *H*^*i*^ positive matrices for *i* = 1. …, *P*.

The algorithm minimizes the objective function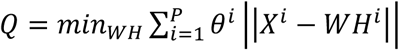.

Once the matrices *W* and *H*^*i*^ *i* = 1, …, *P*have been computed, samples are assigned to clusters based on the W matrix; Each sample is associated with the cluster in which it has the highest weight. The algorithm is implemented into the CRAN R package *intNMF (https://cran.r-project.org/web/packages/IntNMF/index.html)*.

#### 2. Joint and Individual Variation Explained (JIVE)

JIVE^17^ is an extension of PCA to multi-omics data. JIVE decomposes each omics matrix into three structures: a joint factor matrix (J), a omics-specific factor matrix (A) and a residual noise (E):

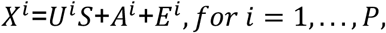

with *E*^*i*^, *A*^*i*^ and *U*^*i*^ are (n_i_ x k) matrices and S is a common score matrix explaining variability across multiple data types.

The algorithm minimizes ‖*E*‖^2^, with *E* ^*i*^ = *X*^*i*^ − *U*^*i*^*S* − *A*^*i*^ and *E* = [*E*^1^… *E*^*P*^]^*T*^.

JIVE is implemented into the R package *r.jive (https://cran.r-project.org/web/packages/r.jive/index.html)*.

#### 3. Multiple co-inertia analysis (MCIA)

MCIA^18^, is an extension of co-inertia analysis (CIA) to more than two omics datasets. MCIA factorizes each omics into omics-specific factors

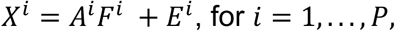

by first applying a dimensionality reduction approach, such as PCA, to each omics matrix *X*^*i*^ separately and then maximizing their co-inertia, i.e. the sum of the squared covariance between scores of each factor:

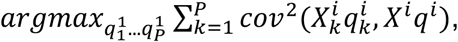

with *var*(*X*^*i*^ *q*^*i*^) = 1and *q*^*i*^ correspond to the global PCA projections. MCIA is implemented in the R package *omicade4 (https://bioconductor.org/packages/release/bioc/html/omicade4.html)*.

#### 4. Regularized Generalized Canonical Correlation Analysis (RGCCA)

RGCCA^21^ is one of the most widely used generalizations of CCA to multi-omics data. Similarly to MCIA, RGCCA factorizes each omics into omics-specific factors:

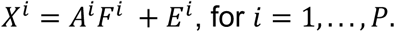

RGCCA maximizes the correlation between the omics-specific factors by finding projection vectors *u*^*i*^ such that the projected data have maximal correlation:

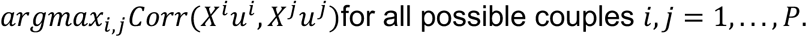

Solving this optimization problem requires inversion of the covariance matrix. However, omics data usually have a higher number of features than samples, and these matrices are therefore not invertible. RGCCA thus apply regularization to CCA. RGCCA is implemented into the CRAN package *RGCCA (https://cran.r-project.org/web/packages/RGCCA/index.html).*

#### 5. iCluster

iCluster^15^ decomposes each omics into the product of a factor matrix that is shared across all omics, and omics-specific weight matrices:

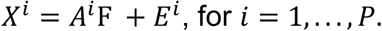

iCluster solves the equation above by first deriving a likelihood-based formulation of the same equation and then applying Expectation-Maximization (EM). The method assumes that both the error *E*^*i*^ and the factor matrix *F* are normally distributed. Finally, clusters are obtained from the factor matrix by applying K-means. The algorithm is implemented into the CRAN package *iCluster (https://rdrr.io/bioc/iClusterPlus/man/iCluster.html)*.

#### 6. Multi-Omics Factor Analysis (MOFA)

MOFA^19^ decomposes each omics into the product of a factor matrix that is shared across all omics, and omics-specific weight matrices:

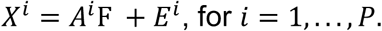

MOFA first formulates the equation above in a probabilistic Bayesian model, placing prior distributions on all unobserved variables *A*^*i*^, *F and E*^*i*^. Then it solves the model by maximizing the Evidence Lower Bound (ELBO), which is equal to the sum of the model evidence and the negative Kullback–Leibler divergence between the true posterior and the variational distribution. MOFA is an extension of Factor Analysis to multi-omics data, but it is also partially related to iCluster. However, differently from iCluster, MOFA does not assume a normal distribution for the errors but supports combinations of different omics-specific error distributions. The code to run MOFA is available at https://github.com/bioFAM/MOFA. The MOFA package further implements an automatic downstream analysis pipeline for the interpretation of the obtained factor and weight matrices through pathways, top-contributing features or percentage of variance-explained interpretation.

#### 7. Tensorial Independent Component Analysis (tICA)

A natural extension of DR methods to multi-omic data is based on the use of tensors, i.e. higher-order matrices. Indeed, all the methods designed for single-omics can be naturally extended to multi-omics with tensors. However, this requires to work with omics data sharing both the same samples and features. Here, to overcome this limitation we ran the tensorial algorithm on the correlation-of-correlation matrix, i.e. the matrix having on rows and columns the samples that are common to all the omics data and having in position (i,j) the correlation of sample i with sample j.

We chose tensorial ICA (tICA)^23^ to represent the tensor-based methods in our benchmark. Considering the multi-omics data organized into a tensor X, the equation solved by tICA is:

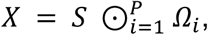

where S is a tensor, with the same dimension of X, and composed of *S*_1_… *S*_*P*_ random variables mutually statistically independent and satisfying *E*[*S*_1_… *S*_*P*_] = 0 and *Var*[*S*_1_… *S*_*P*_] = *I*and ⊙ denotes the tensor contraction operator.

Thus, tICA searches for independent signals. Since biological processes are generally non-Gaussian and often sparse, the assumption of tICA can improve the deconvolution of complex mixtures and hence better identify biological functions and pathways underlying the multi-omics data. Given that multiple tensorial versions of ICA exist, we considered the tensorial fourth-order blind identification (tFOBI), whose implementation in R is available in ^23^.

#### 8. Multi-Study Factor Analysis (MSFA)

MSFA is a generalization of Factor Analysis (FA), which models the omics matrices *X*^*i*^ as the sum of data-specific and shared factors:

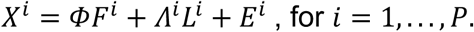

where*E*^*i*^ has a multivariate normal distribution and the marginal distributions of *F*^*i*^,*L*^*i*^ and *X*^*i*^ are multivariate normal. MFSA is implemented in R and available at https://github.com/rdevito/MSFA.

#### 9. Data fusion (scikit-fusion)

The data-fusion approach (scikit-fusion)^22^ is based on two steps. First, two groups of matrices are constructed from the multi-omics data: relation (*R*) and constraint (*C*) matrices. The R matrix encodes relations inferred between features of different omics (e.g., genes to proteins) and the matrix C describes relations between features of the same omics (e.g. protein-protein interactions). The matrix C thus corresponds to the side information that scikit-fusion can consider in the factorization. Then, tri-matrix factorization is used to simultaneously factorize the various relation matrices R under constraints C. Given that the R and C matrices are block-matrices, with element *R*_*ij*_containing a relation between the elements of the i-th omics and those of the j-th, the matrix tri-factorization is applied separately to each block:

*R*_*ij*_≈ *G*_*i*_*S*_*ij*_*G*_*j*_, with *G*_*i*_shared across all the *R*_*ip*_ for *p* = 1… *P*(matrices that relate the i-th object to others).

Hence, scikit-fusion can naturally combine additional side-information in the factorization of the multi-omics data, such as protein-protein interactions, Gene Ontology annotations. It is implemented in Python and available at https://github.com/marinkaz/scikit-fusion.

### Factor selection for performance comparisons

The jDR approaches make different assumptions on the cross-omics constraints of the factors. The various jDR can be thus classified in shared factors, omics-specific factors and mixed factors approach. To use the factor matrices to compare the performances of the various jDR methods, e.g. to cluster the samples based on the factors, we had to select which factor matrix to use for each jDR. Shared factors jDR methods compute a unique factor matrix, which is used in our benchmark. Omics-specific jDR methods compute a factor matrix for each omics dataset. In these cases, we selected the factor matrix associated with transcriptomic data for our benchmark. However, jDRs with omics-specific factors maximize correlation or co-inertia between the various omics-specific factor matrices. The values of the transcriptomic factor matrix are then influenced by the other omics. Finally, for mixed factors jDRs methods, we considered the joint factor matrix *F*. As a consequence, all jDR methods with omics-specific and mixed factors contain in their factorization more information than those considered here for sake of comparison.

### Data simulation

The simulated multi-omics datasets have been produced by the *InterSIM* ^24^ CRAN package. *InterSIM* simulates multiple interrelated data types with realistic intra- and inter-relationships based on the DNA methylation, mRNA gene expression, and protein expression from TCGA ovarian cancer data. We generated 100 simulated datasets, with a number of clusters set by the user. We considered five, ten and fifteen clusters in this study. The proportion of samples belonging to each subtype is also set by the user, while we considered here two conditions with equally sized clusters and variable random sizes, respectively.

### Clustering of factor matrix

To identify the clusters of samples starting from the jDR factor matrix, we applied k-means clustering to the factor matrix. We chose k-means for clustering in agreement with the use of k-means in iCluster and euclidean distance in intNMF for clustering. As k-means clustering is stochastic ^25^, we performed clustering 1000 times and computed a consensus consisting in the most frequent associations between samples and clusters

### Comparing jDR algorithm clusters to ground-truth clusters

The matching between the ground-truth clustering and the clustering inferred by the various jDR algorithms is measured with the Jaccard Index (JI). JI is a similarity coefficient between two finite sets A and B, defined by the size of the intersection of the sets, divided by the size of their union: 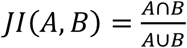. It takes its values in [0;1].

### Selection of the clinical annotations

The clinical annotations selected for benchmark testing are “age of patients”, “days to new tumor”, “gender” and “neo-adjuvant therapy somministration”. This set of annotations is obtained after excluding redundant annotations (e.g. “age_at_initial_pathologic_diagnosis” and “years_of_initial_pathologic_diagnosis”), annotations having missing values for more than half of the samples, and annotations having no biological relevance (e.g. “vial_number”, “patient_id”). Four clinical annotations are available for nine or ten out of ten cancer types, while the others are only present for six or fewer cancer types (with most of them being available only for one or two cancer types).

### Selectivity score

We define the selectivity as:

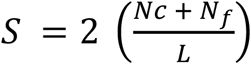

where Nc is the total number of clinical annotations associated with at least a factor, N_f_ the total number of factors associated with at least a clinical annotation, and L the total number of associations between clinical annotations and factors. S has a maximum value of 1 when each factor is associated with one and only one clinical/biological annotation, and a minimum of 0 in the opposite case. An optimal method should thus maximize its number of factors associated with clinical/biological annotations without having a too low selectivity.

### Testing the biological enrichment of metagenes

To test if metagenes are enriched in biological annotations, we used *PrerankedGSEA*, implemented in the *fgsea* R package. In *prerankedGSEA*, each metagene is considered as a ranking of genes, and the significance of the association of a biological annotation with the higher or lower part of the ranking is tested. We considered as biological annotations Reactome pathways, Gene Ontology (GO) and cancer Hallmarks, all obtained from MsigDB^30,31^.

### Quality of single-cell clusters

To evaluate the quality of the clusters obtained from single-cell multi-omics data, we employed the C-index measure^32^, an internal clustering evaluation index comparing the distance between intra-cluster points and inter-clusters points. The C-index has values in [0,1] and should be minimum in an optimal clustering.

---

**Supp Table 1.**
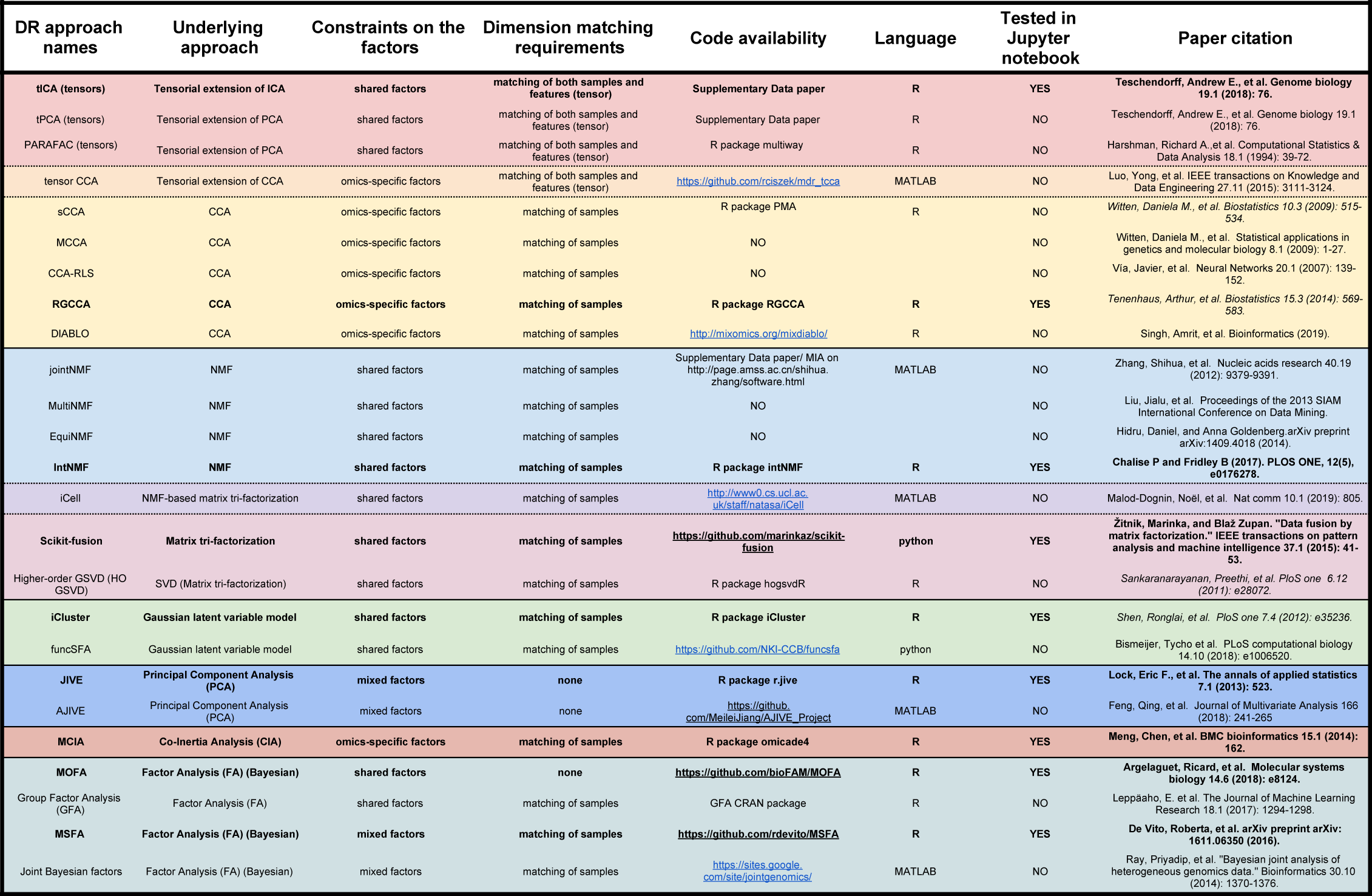
Extended list of existing DR multi-omics integrative algorithms. The algorithms are grouped based on their underlying approach. The columns of the table report, the names of the DR method, its underlying approach, the contrained that it assumes on the factors, if it requires to match features and/or samples, the link to the code, the language of the code, if the algorithm has been tested in our benchmark and the link to the paper of the method.

**Supp Figure1.**
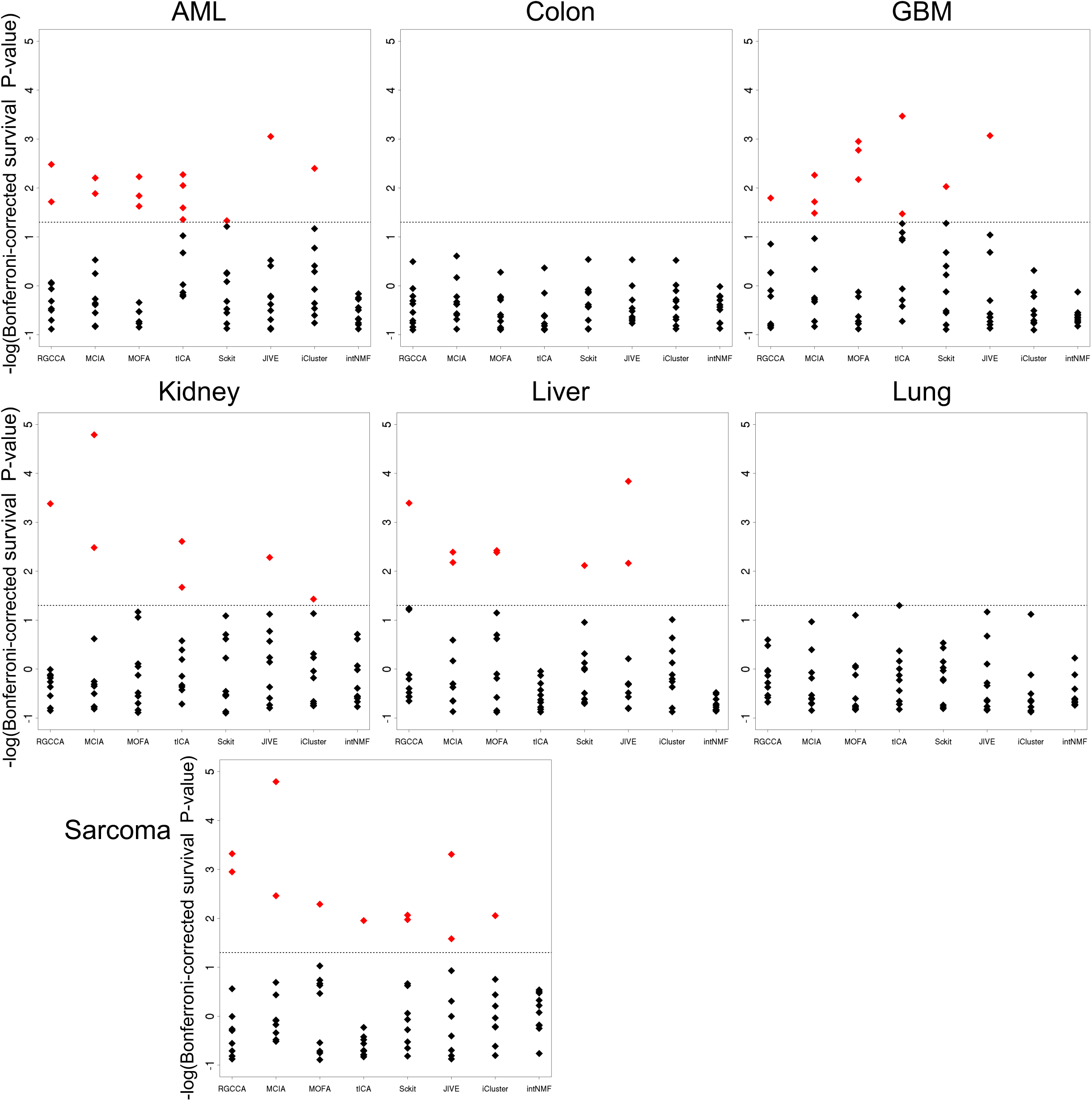
Identification of factors predictive of survival in cancer samples by the jDR methods. For each method the Bonferroni-corrected p-values associating each of the 10 factors to survival (Cox regression-based survival analysis) are reported. The dot lines correspond to a corrected p-value threshold of 0.05.

**Supp Figure 2.**
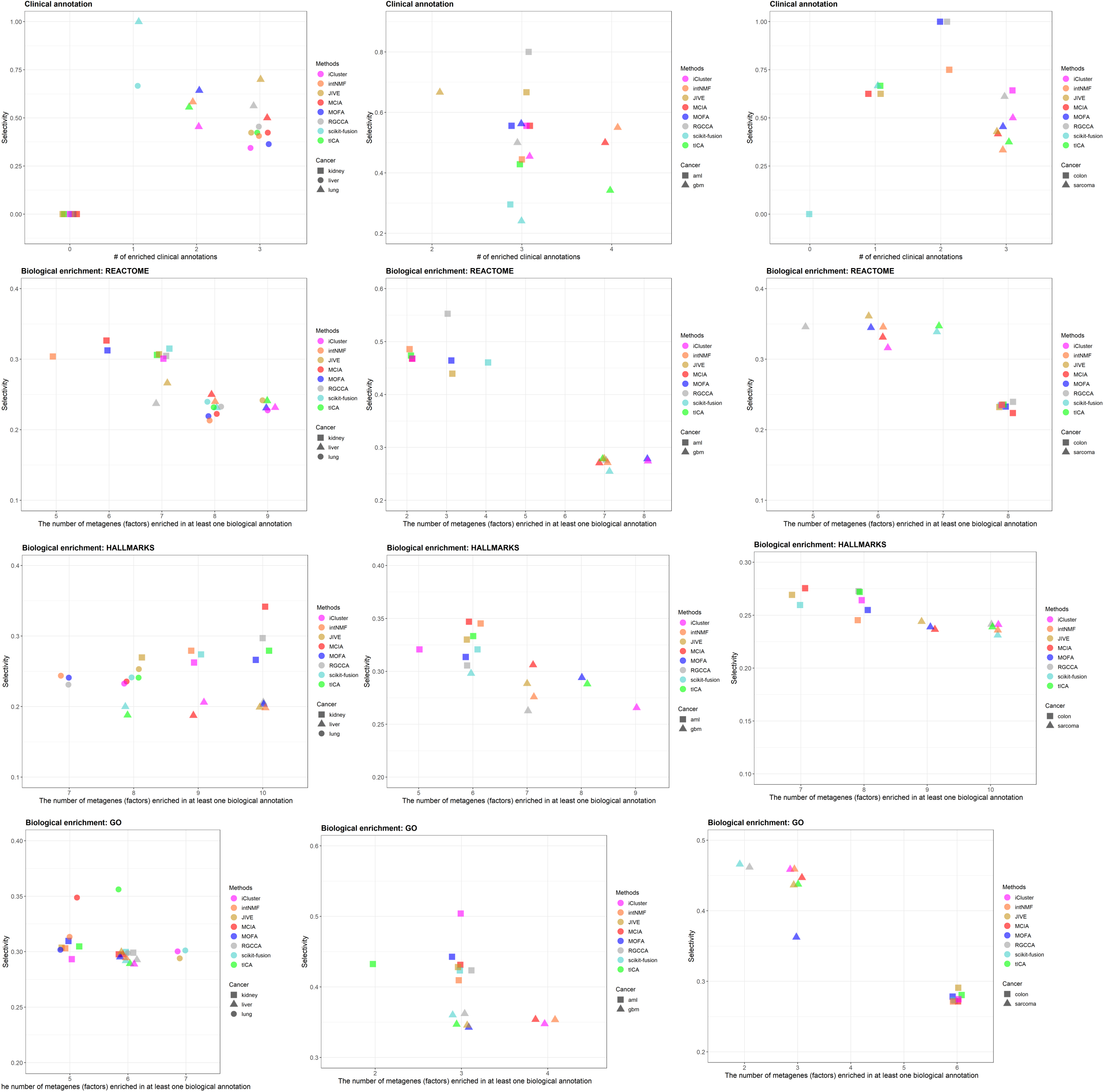
Identification of factors associated with clinical annotations, and metagenes associated with biological annotations in cancer samples, by the jDR methods. For clinical annotations, the plot represents, for each method, the number of clinical annotations enriched in at least one factor together with the selectivity of the associations between the factors and the clinical annotations (Method). For the three annotation sources (MsigDB Hallmarks, REACTOME and Gene Ontology), the number of metagenes identified by the different jDR methods enriched in at least a biological annotation are plotted against the selectivity of the associations between the metagene and the annotation.

